# Chiral and nematic phases of flexible active filaments

**DOI:** 10.1101/2022.12.15.520425

**Authors:** Zuzana Dunajova, Batirtze Prats Mateu, Philipp Radler, Keesiang Lim, Philipp Velicky, Johann Georg Danzl, Richard W. Wong, Jens Elgeti, Edouard Hannezo, Martin Loose

## Abstract

The emergence of large-scale order in self-organized systems relies on local interactions between individual components. During bacterial cell division, the tubulin-homolog FtsZ polymerizes into treadmilling filaments that further assemble into a cytoskeletal ring. Although minimal *in vitro* assays have shown the striking self-organization capacity of FtsZ filaments, such as dynamic chiral assemblies, how these large-scale structures emerge and relate to individual filament properties remains poorly understood. To understand this quantitatively, we combined minimal chiral active matter simulations with biochemical reconstitution experiments. Using STED and TIRF microscopy as well as high-speed AFM, we imaged the behavior of FtsZ filaments on different spatial scales. Simulations and experiments revealed that filament density and flexibility define the local and global order of the system: At intermediate densities, flexible filaments organize into chiral rings and polar bands, while an effectively nematic organization dominates for high filament densities and for mutant filaments with increased rigidity. Our predicted phase diagram captured these features quantitatively, demonstrating how filament flexibility, density and chirality cooperate with activity to give rise to a large repertoire of collective behaviors. These properties are likely important for the dynamic organization of soft chiral matter, including that of treadmilling FtsZ filaments during bacterial cell division.

## Introduction

In active systems, the emergence of large-scale order relies on a combination of local interactions between components and microscopic energy consumption. One typical property of such self-organizing systems is spontaneous motility of their constituents. For example, cytoskeletal filaments can be transported due to the activity of motor proteins or move due to treadmilling polymerization dynamics. Dynamic interactions between active constituents can lead to complex collective behavior and phases not found at equilibrium, which have been under intense experimental and theoretical investigation in the past decade^1–7^. Reconstituted mixtures of actin or microtubule filaments with motor proteins self-organize into moving swarms, vortices, and travelling waves^1,5,8^. In living systems, such emergent behaviors can underlie a wealth of key biological phenomena such as single and collective cell motility^9,10^, cell division^11^ and organism morphogenesis^12,13^.

Active matter systems can be classified according to the symmetry of its constituents (e.g. polar or nematic)^14–16^. In particular, chiral active matter has recently attracted attention, where the constituents are either asymmetric in shape or perform a circular self-propelled motion. This includes curved cytoskeletal filaments, asymmetric synthetic swimmers or cell types displaying chiral motions on 2D substrates^17–19^. However, how to relate the microscopic properties of active constituents to the large-scale outputs relevant for biology remains an outstanding challenge in the field^16^. For instance, biological settings often involve highly dense systems with attraction, as well as deformable constituents which can change their shape as a function of external forces or crowding^6^. How flexible constituents self-organize and how their local deformability contributes to large-scale collective features remain poorly understood, both theoretically and experimentally.

One example of treadmilling filaments are polymers formed by the protein FtsZ, a tubulin homolog that organizes cell division in almost all bacterial, and some archaeal, species^20,21^. FtsZ forms single stranded filaments that grow from one end and shrink from the opposite end^22^, driven by the hydrolysis of guanosine triphosphate (GTP). FtsZ filaments can also interact laterally, which facilitates the condensation of filaments from a diffuse organization into a tight ring-like structure called the Z-ring^23–26^. The density of this ring further increases during constriction until the cell is split into two^23,27^. FtsZ also forms treadmilling filaments *in vitro*, which organize into cytoskeletal patterns of moving bands and chirally rotating rings^28^. While lateral interactions between treadmilling filaments play an important role for Z-ring assembly in living cells and the emergence of cytoskeletal structures *in vitro*, the physical properties and interaction rules of treadmilling filaments on a membrane surface are currently not well known. Here, by studying the self-organization of FtsZ filaments at different spatial and temporal scales, we elucidate the quantitative principles that govern the emergence of different collective cytoskeletal organizations from the local interactions of active constituents.

### Modes of FtsZ filaments self-organization at different densities

To understand the properties of FtsZ filament bundling, we first explored experimentally the phase space of possible organizations of FtsZ filaments as a function of their density. We used a previously established *in vitro* reconstitution assay, where treadmilling FtsZ filaments are recruited to the surface of a supported bilayer by the membrane anchor FtsA^28^. Using total internal reflection fluorescence (TIRF) microscopy, we could record the emergent behavior of the membrane-bound filaments. We found that we could control the density of membrane-bound FtsZ filaments by changing the FtsZ concentration in the buffer solution. When we increased the FtsZ bulk concentration from around 0.6 to 5 µM while keeping the FtsA concentration constant, the fluorescence intensity of FtsZ on the membrane increased linearly until it saturated at concentrations higher than 3 µM (**Supplementary Fig. 1a**). At the same time, we found that the large-scale organization of the filaments changed with their density (**Fig. 1a, Supplementary Movie 1**).

**Figure 1:**
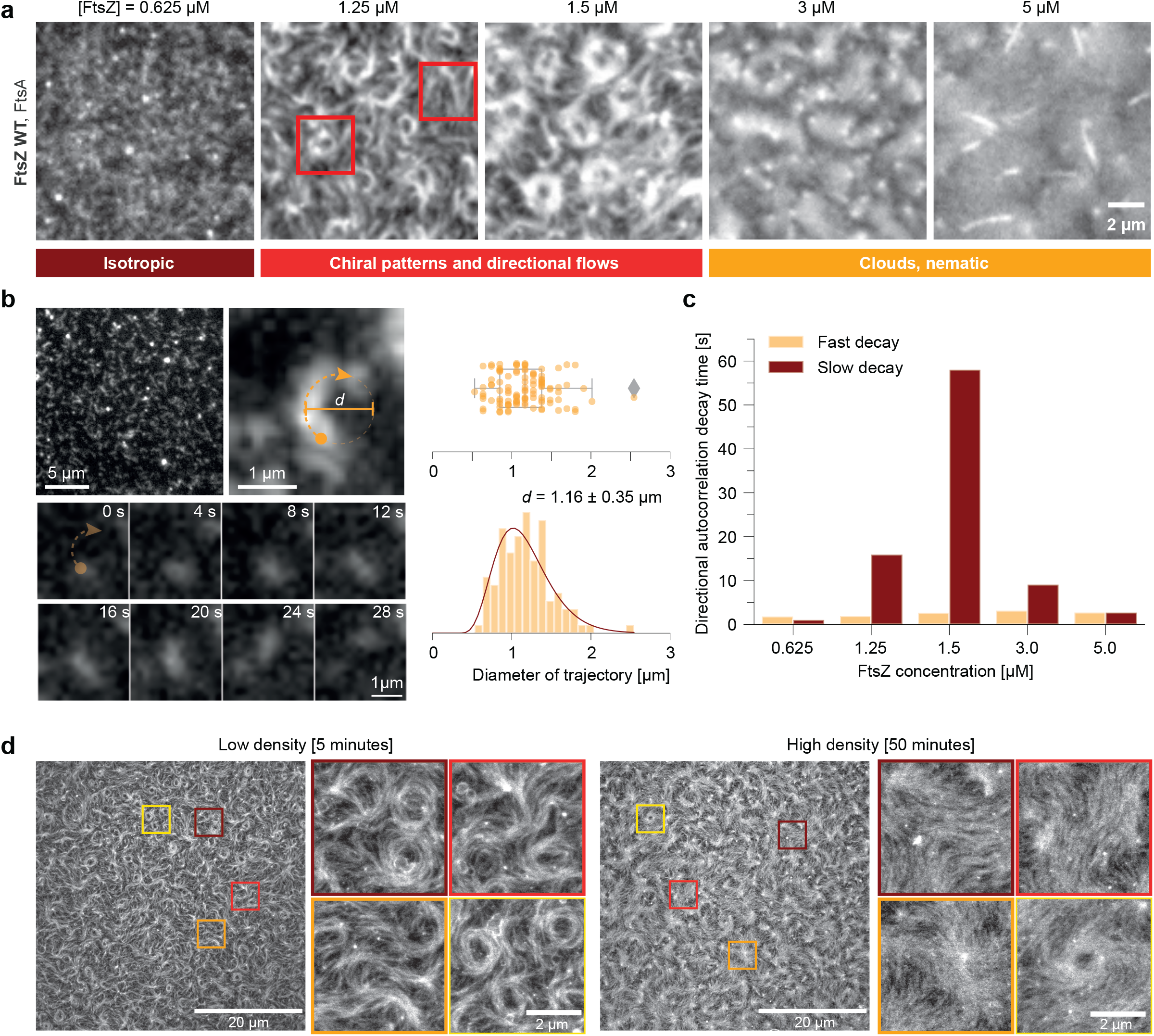
The orientation of FtsZ filament bundles changes with increasing density on SLBs. **a**, Representative TIRF micrographs of Alexa488-FtsZ at increasing FtsZ and constant FtsA concentrations. Below 0.625 µM FtsZ, filaments do not form higher order structures. At 1.25 and 1.5 µM, FtsZ forms rotating rings and directionally moving filament bundles. This organization is lost at 3 µM and 5 µM FtsZ, at which filaments densely cover the membrane surface. Scale bars are 2µm. **b**, Representative images of a trajectory of a single FtsZ filament at 0.5 µM (left) and distributions of measured curvatures. The filament moves along a curved path, corresponding to an apparent diameter of 1.15 ± 0.346 µm (std, n = 105). Scale bars are 5 and 1 µm. **c**, Decay constants from fitting a bi-exponential function to directional autocorrelation curves from treadmilling trajectories. For intermediate FtsZ concentrations, the curves are best fitted assuming a fast and slow decay constant, consistent with persistent directional flows. **d**, Representative STED micrographs of 1.5 µM Atto633-FtsZ tethered to SLBs by 0.2 µM FtsA at low (5 minutes after starting the experiment, top) and high (50 minutes, bottom) densities. First, rotating rings and moving bundles first coexisted on the membrane surface. With increasing filament density ring-like structures disappeared. Scale bars: 20 or 2 µm.

At FtsZ concentrations lower than 0.6 µM, individual filaments traveled across the membrane surface^28^. These filaments were too short to measure their intrinsic curvature by fluorescence microscopy, but maximum intensity projections of time lapse movies revealed that their trajectories followed a curved path corresponding to a circle with a diameter of 1.16 µm ± 0.35 µm and heavily biased in the clockwise direction (**Fig. 1b, Supplementary Fig. 1b**).

When we increased the FtsZ bulk concentrations to 1.25 and 1.5 µM, FtsZ filaments organized into chiral rotating rings that persisted for around 5-6 min and coexisted with comet-like structures and moving bundles of treadmilling filaments as observed previously^28^. When we calculated the directional autocorrelation of cytoskeletal flows, we found a long decay time of 16 and 68 seconds at 1.25 and 1.5 µM FtsZ respectively, consistent with persistent, long-range polar motion of filament bundles. In fluorescence recovery after photobleaching experiments we found that the mean lifetime of FtsZ monomers in filament bundles was only around 7 s, suggesting that their directional motion is a collective property of the system (**Fig. 1c, Supplementary Fig. 1c**).

At bulk concentrations of 3 µM and higher, filaments densely covered the membrane surface without apparent large-scale organization or directional movements. FtsZ filaments still continuously exchanged monomers at these high densities, with a recovery half-time of around 15s (**Supplementary Fig. 1d**).

These features were also recapitulated using time-lapse stimulated emission depletion (STED) microscopy. Compared to TIRF microscopy, STED provided higher resolution but did not allow for long-term imaging of the filament pattern. It still showed that rings form preferentially at intermediate densities, and are comprised of many transiently interacting FtsZ filaments (**Fig. 1d, Supplementary Movie 2, Supplementary Fig. 1e)**. The apparent weak interactions between filaments suggest that their local polar orientation is an emergent property of the ensemble, rather than the result of highly specific static interactions, such as residue contacts between individual filaments (**Supplementary Fig. 1f, g**).

Together, these observations and quantifications show that there are density-dependent transitions of the large-scale organization and motion of treadmilling filaments.

### Collective filament organization as a function of bending rigidity and attraction

Given our experimental findings that single FtsZ filaments displace along curved, chiral paths, and previous theoretical work showing that non-adhesive chiral self-propelled filaments could organize into ring-like phases at intermediate densities^2^, we explored whether a minimal coarse-grained model could quantitatively reproduce the observed phenomenology. To explore the limits of strong adhesion, high densities and low bending rigidity, we modeled this system on a mesoscopic level by a collection of overdamped self-propelled semi-flexible filament bundles in two dimensions^29,30^ (see section Material & Methods for more details on the simulation framework). Each semi-flexible filament bundle is simulated as a worm-like chain, with bending rigidity described by the potential V*bend*, and is self-propelled with tangential force *Fip* to describe an effective treadmilling velocity v0= *Fip* /*γ*, where *γ* is the friction coefficient with the membrane. In the following, we use the flexure number^29^ *ℱ*, defined as the ratio of self-propulsion forces to bending rigidity 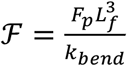, as a measure of filament flexibility (inverse of rigidity). To account for the observed chirality, filaments are considered to have spontaneous signed curvature with rest angle *θ*_0_. An effective thermal noise force 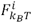 accounts for the many sources of noise in the system (**Fig. 2a**). Finally, based on previous studies of FtsZ filament-filament interactions^26,32,33^ and our own observations (**Fig. 1d**), we considered mid-range attractive interactions between filaments (*Vpair*).

**Figure 2:**
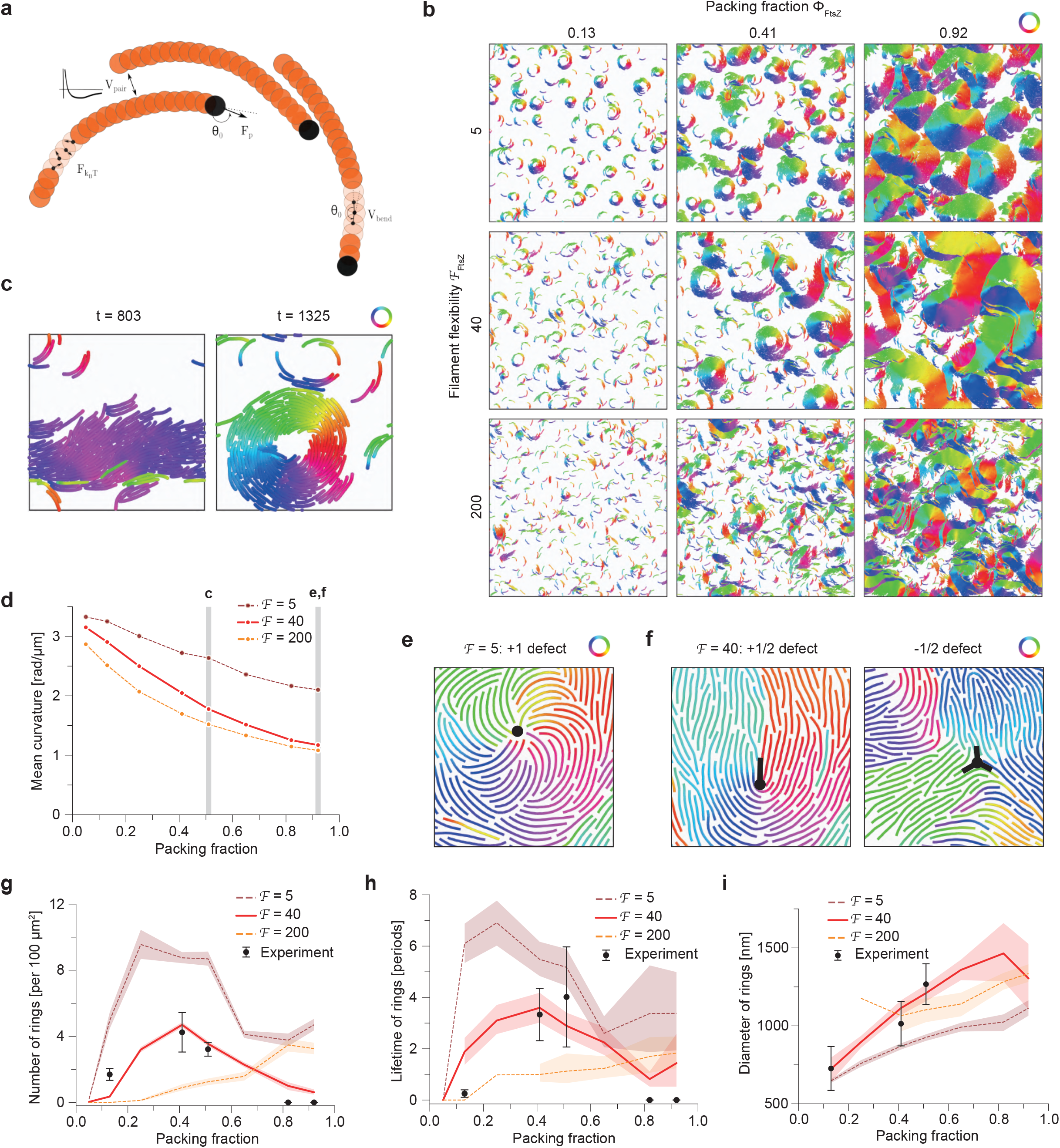
Numerical simulation of FtsZ WT self-organization. **a**, Scheme of the simulation model. **b**, Phase diagram of the large-scale patterns (*L* = 212 8) with varying filament flexibility (measured by flexure number *ℱ*, vertical axis) and density (horizontal axis). Filaments are color-coded according to the orientation of the bond vectors between beads. We observe ring-like self-organization of rigid filaments (*ℱ* = 5), spatial coexistence of chiral rings and polar bands in regime of semiflexible filaments (*ℱ* = 40) and disordered patterns with flexible filaments (*ℱ* = 200). **c**, Temporal coexistence of chiral rings and polar bands in a small simulated system (*L* = 42 8) of intermediate density (*Φ* = 0.5) and filament flexibility (*ℱ* = 40). Filaments are color-coded according to the orientation of the bond vectors between beads. **d**, Average filament curvature with varying density, showing density-driven straightening. Three values of filament flexibility are shown as in the phase diagram in b. **e**, Spiral defect for rigid filaments (*ℱ* = 5) and high density (*Φ* = 0.9). Only bonds of the filaments (without the full diameter of beads) are presented for clarity. Filaments are color-coded according to the orientation of the bond vectors between beads. **f**, Nematic defects in high density (*Φ* = 0.9) of semiflexible filaments (*ℱ* = 40). Only bonds of the filaments (without the full diameter of beads) are presented for clarity. Filaments are color-coded according to the orientation of the bond vectors between beads. **g-i**, Quantitative comparison of ring density, lifetime and diameter between simulations and experiments (see Methods for details on the quantification and comparisons). Red solid line corresponds to the best fit of filament flexibility (*ℱ* = 40).

We observed that the filament self-organization was markedly different with varying filament flexibility (**Fig. 2b, Supplementary Fig. 2a, Supplementary Movie 3**). In the case of very rigid, curved filaments, ring-like patterns and vortices dominated the system throughout the experimentally explored density range. However, this was not the case for semiflexible filaments, which displayed two density-driven transitions as seen in experiments. Moreover, in this regime we could observe a co-existence, both temporal and spatial, of chiral rings and nematic-like, straighter traveling bands characterized by low spontaneous curvature. Interestingly, this was in strong qualitative agreement with our experimental observations, where filaments can adopt a wide range of curvatures even at intermediate densities^34^ and where rings are interspersed with less ordered filament assemblies **(Supplementary Fig. 1e)**. In addition, as in our experimental data, we observed dynamical interconversion between ring and band patterns (**Fig. 2c, Supplementary Fig. 2b, Supplementary Movie 4**). At the other extreme, filaments with very low bending rigidity deformed too easily and formed only rare and unstable chiral rings at all density tested. Together, this phase diagram argues that filament flexibility could be a key parameter for FtsZ self-organization.

More quantitatively, by analyzing the average filament curvature in our simulations, we found that the nature of self-organization in high densities was markedly different with varying filament flexibility (**Fig. 2b, d**). In the case of very rigid filaments, even above the critical density where well-defined rings disappeared in simulations, we found that individual filaments were still highly curved and could self-organize into spiral patterns. These were characterized by the presence of +1 topological defects and displayed chiral rotation dynamics (**Fig. 2e)**. However, semiflexible filaments displayed pronounced decrease in their average curvature with increasing density, so that they formed effectively an active nematic phase, characterized by spontaneous appearance and movement of +1/2 and -1/2 topological defects (**Fig. 2f, Supplementary Movie 5**).

Before turning to a more quantitative assessment of the match between data and modelling, we also investigated the effect of attraction and noise (Peclet number) for the phase diagram. For low attraction, we systematically observed a transition from disordered patterns to rings above a critical density, as previously reported^2^. Interestingly however, for strong FtsZ attraction, this transition was largely lost, with rings able to form even at the lowest experimental densities (**Supplementary Fig. 2c, 3a**), similarly to the case of very rigid filaments. From a physical perspective, this is due to ring formation being energetically favored, instead of being an active kinetic state in the case of purely repulsive self-propelled filaments. The Peclet number also strongly affected the first transition, as well as the overall density of rings in the system (**Supplementary Fig. 2d, 3b**). Above a second threshold of density, we could observe in all cases a loss of ring patterns.

### Quantitative comparison between model and experiments

To more systematically and quantitatively compare simulations and experiments, we first sought to constrain model parameters. From our observations of single-filament trajectories (**Fig. 1b**) and previously published values^28,35^ we considered a treadmilling speed of *v0* = 0.04 5m/s, and estimated the packing fraction of filaments 6 based on our calibration experiments (**Supplementary Fig. 1a)**. The aspect ratio of filaments in our mesoscopic simulations corresponded to bundles with thickness of 5 FtsZ filaments of length 400 nm (see section Material & Methods for more details on parameter estimation). Estimation of filament chirality based on curvature of single-filament trajectories was noisy and limited by the resolution of fluorescence microscopy, but still allowed us to guide parameter space by estimating *θ*_0_ to yield the single-filament rotation diameter of 500-1000 nm, as well as a lower bound for the Peclet number of ∼100 (**Supplementary Fig. 3k**). Therefore, after this parameter estimation, we were left with only three fitting parameters, which we systematically explored in our simulations: the adhesion strength *ε*, the filament flexibility measured by flexure number *ℱ* and noise strength measured by Peclet number *Pe*.

Interestingly, intermediate values of filament flexibility, as well as low to intermediate attraction, provided a good match for a number of qualitative and quantitative features of our dataset. Firstly, we could quantitatively reproduce not only the two thresholds of appearance and disappearance of rings as a function of density, but also the absolute probability of ring formation in the intermediary density region (**Fig. 2g**). Strikingly, when measuring ring life-time (normalized by period of filament rotation) in both experiments and simulations, we found excellent agreement with our predicted parameter regime (**Fig. 2h, Supplementary Fig. 3c, d**). Finally, our simulations predicted that the diameter and thickness of rings should monotonously increase with density, a feature in agreement with the data (**Fig. 2i, Supplementary Fig. 3e, f**). Overall, this demonstrates that the transitions seen in the data as a function of density can be quantitatively explained by a simple theoretical framework of flexible chiral active filaments. Interestingly, this analysis suggests FtsZ filaments being much more flexible than previously anticipated^2,30^. This prompted us to test more directly how FtsZ filament conformation changes at the microscopic level as a function of increasing density.

### High-speed-AFM imaging of dynamic FtsZ filaments

As fluorescence microscopy was not sufficient to visualize individual filaments inside of dense filament bundles, we applied high speed atomic force microscopy (HS-AFM) on treadmilling, membrane-bound filaments, which allowed temporal imaging of biological samples at high lateral resolution down to 2-3 nm and at up to 20 frames per second^36,37^. We were able to image individual filaments that dynamically moved across the membrane surface and increased in density during the course of the experiments. These filaments rapidly moved in and out of the field of view, which made it difficult to track individual filaments over time. However, these time-lapse movies confirmed that neighboring filaments did not form stable lateral contacts; instead they displayed fast lateral fluctuations on the membrane surface (**Fig. 3 a, Supplementary Movie 6**). This suggests that the attraction energy was comparable or smaller to fluctuations, consistent with our numerical simulations. We could also reconstruct filament shape as a function of density. Interestingly, we found a gradual decrease in filament curvature as a function of overall surface density, which fitted well our theoretical prediction for semiflexible filaments (**Fig. 3b**). At the highest densities, filaments had no discernable intrinsic curvature and showed hallmarks of active nematics, such as spontaneous formation of +1/2 and -1/2 topological defects (**Fig. 3c**), another feature generically observed in our simulations (**Fig. 2f**). This confirms that their organization at high densities is governed by nematic rather than polar filament-filament interactions.

**Figure 3:**
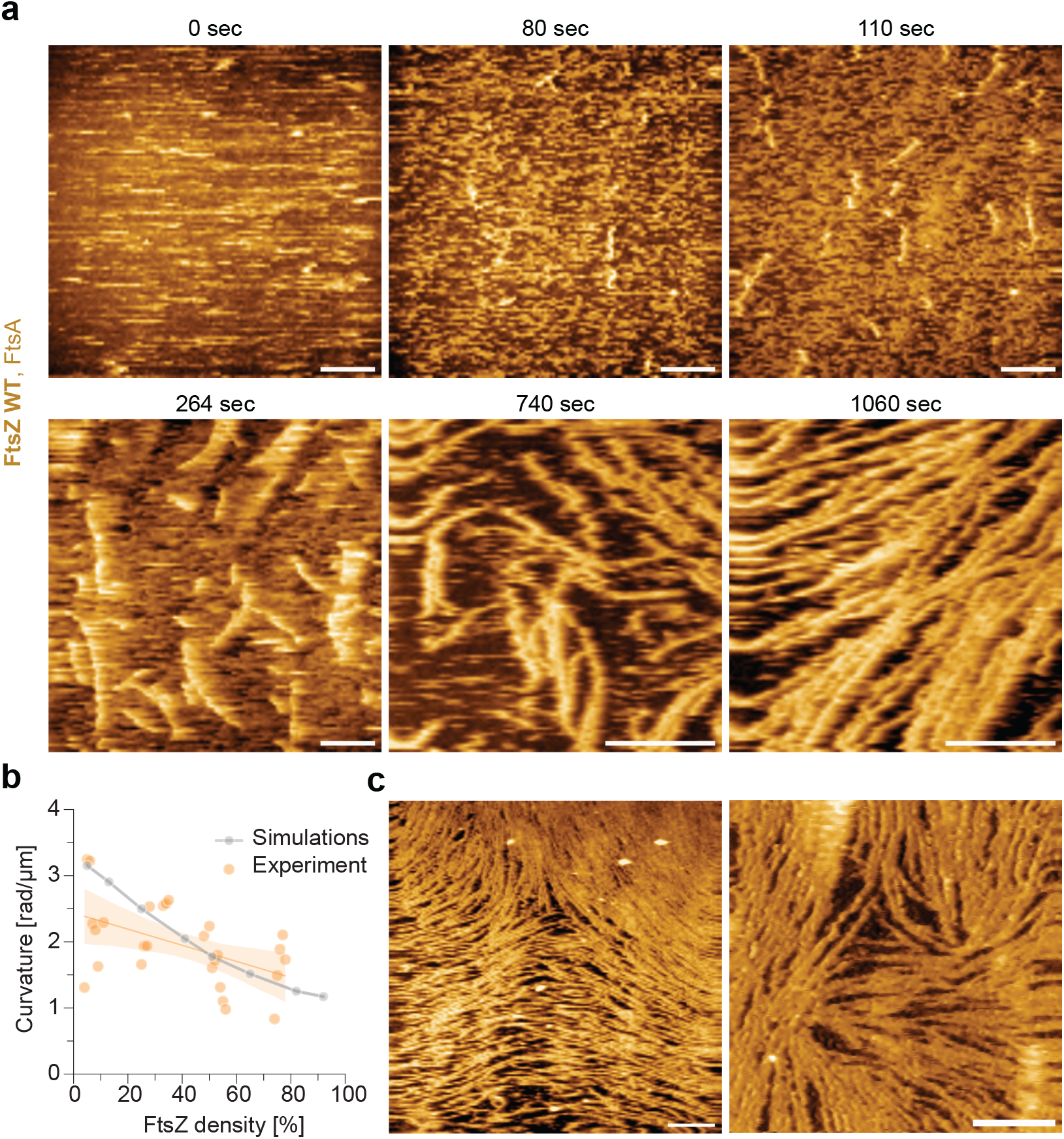
High-speed AFM imaging shows nematic organization at high densities of FtsZ filaments. **a**, Representative HS-AFM time-lapse experiment, showing an increase in density of FtsZ WT filaments with passing time. The filaments move less and become more stable with increasing time. Scale bars are 500 nm. **b**, Curvature of FtsZ WT filaments at increasing densities from experiments (orange) and simulations (gray). With increasing densities, the curvature of individual filaments decreases. **c**, At high densities FtsZ WT filaments show nematic order with topological defects ([FtsZ] = 1.5 µM, left and 4.0 µM, right). Scale bars: 500 nm.

### Testing the model by changing filament conformation

Since our theory suggests that semiflexible filaments and intrinsic chirality are crucial to explain the density-dependent transitions that we observed experimentally, we sought to challenge it further by using constituents with altered properties, i.e. straight filaments with enhanced bending rigidity. We reviewed previously described FtsZ mutants and identified FtsZ L169R as an interesting candidate protein. This mutant was previously described to have increased lateral interactions, as it shows enhanced bundling *in vitro* and *in vivo*^38,39^. We used Alphafold^40,41^ predictions of a FtsZ-FtsZ dimer using the crystal structure of FtsZ filaments from *Staphylococcus aureus* as a template^42^ and found that the positively charged Arginine residue at position 169 is in fact located at the longitudinal interface, facing towards a negatively charged Glutamic acid residue at position 276 of the neighboring monomer (**Fig. 4a**). Accordingly, the L169R mutation could result in an additional salt-bridge between two monomers in the filament, which could straighten and stiffen the resulting filaments. Indeed, our HS-AFM experiments with this FtsZ mutant confirmed our hypothesis and also revealed a number of key differences from the wildtype protein. First, while wildtype filaments moved in and out of the field of view, filaments of FtsZ L169R mutant protein appeared to be more static. This loss of mobility suggests that their kinetic polarity is perturbed (**Supplementary Movie 7**). We indeed found that many filaments displayed bidirectional growth and sudden shrinkage events rather than treadmilling behavior. Second, our HS-AFM experiments showed that single filaments appeared to have lower intrinsic curvature and less fluctuations than filaments of the wildtype protein even at very low densities (**Fig. 4b**). When we analyzed the shape of individual filaments at different densities, we found that the curvature of wildtype filaments decreased two-fold with increasing density, while mutant filaments were almost straight at all densities studied (**Fig. 4c**). At the same time the persistence and contour lengths were about two times higher for FtsZ L169R filaments and further increased with their densities (**Fig. 4d, e**). We also found that mutant filaments showed transient filament interactions and height profile similar to the wildtype filaments, but with a two times smaller mean filament distance (**Supplementary Fig. 4a-c**)^38^.

**Figure 4:**
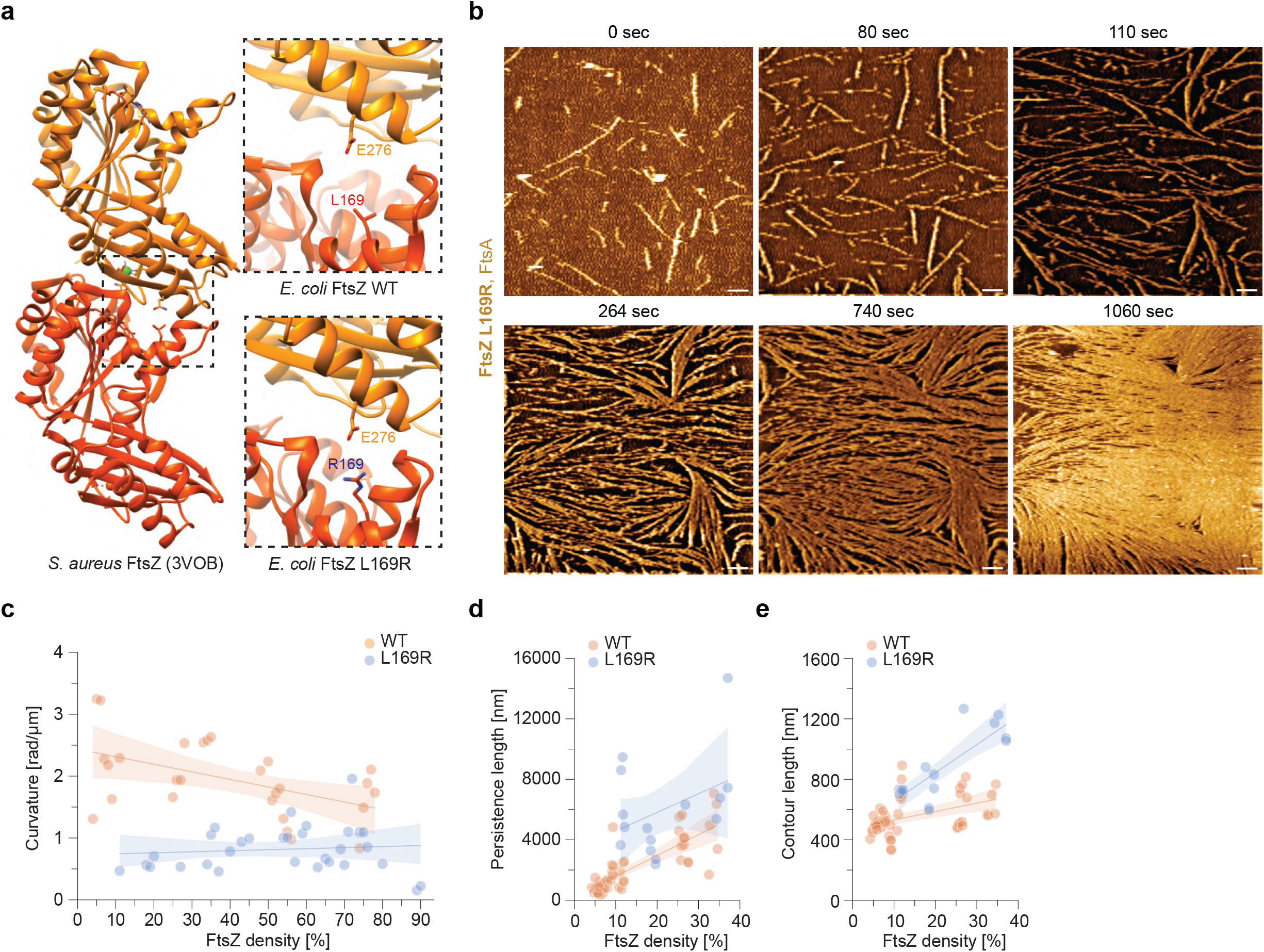
A point mutation in FtsZ L169R changes the properties of FtsZ filaments. **a**, Ribbon model of the *Staphylococcus aureus* FtsZ filament (PDB: 3VOB), left, and the longitudinal interface predicted for *E. coli* FtsZ WT (right, top) and L169R (right, bottom). The Leucine to Arginine mutation likely enables a novel salt bridge, which stabilizing the FtsZ filament. **b**, Representative HS-AFM time-lapse experiment, showing an increase in density of FtsZ L169R filaments. Already at low densities the mutant filaments are less dynamic and more rigid. Scale bars are 500nm. **c**, Curvature of FtsZ L169R filaments as a function of density, showing that it is lower than of FtsZ WT filaments at all tested densities. **d**, Persistence length of FtsZ L169R filaments as a function of density, showing a two times higher value compared to FtsZ WT. **e**, Contour length of FtsZ L169R as a function of density, increasing faster than for FtsZ L169R. Data shown in **d, e** is taken from experiments at low densities (<40%), as FtsZ L169R filaments at higher densities can be longer than the field of view and limiting quantification.

Given that we previously had assumed a local polar order between filaments in our mesoscopic bundle simulations, we now wanted to test the effect of the filament properties of FtsZ L169R. We thus quantified local filament-filament alignment in toy-simulations with different filament parameters. Interestingly, we found that all changes observed in the FtsZ L169R mutant (longer, less chiral, or more persistent filaments with slower treadmilling) went in the same direction of decreasing polar filament alignment **(Supplementary Fig. 4e-h)**. Combining these three experimentally observed static properties thus predicted a strong decrease of local polarity sorting (**Fig. 5a, b, Supplementary Fig. 4d, Supplementary Movie 8**). We then examined the experimental large-scale patterns of mutant FtsZ filaments via TIRF microscopy. FtsZ L169R polymerized into a cytoskeletal network of dynamic filament bundles at concentrations between 0.6 µM and 3 µM (**Fig. 5c, Supplementary Movie 9**). We found that while the mutant reached saturation on membranes with FtsA similarly to the wildtype, its turnover was 2-3x slower and that it hydrolyzed GTP 3x slower compared to the wildtype (**Supplementary Fig. 4i-k**). Furthermore, we did not observe any directional flows or chirally rotating rings on the membrane surface. In addition, differential imaging did not generate directional moving speckles, confirming that local polar arrangement of filaments is strongly impaired (**Supplementary Fig. 4l**)^35^. These features were well-recapitulated by the same large-scale bundle simulations as before, but with the HS-AFM-derived properties for the mutant filaments: longer, less chiral and more persistent filaments with slower kinetics *Fp* (**Fig. 5d, Supplementary Movie 10**). Altogether, this shows that our simplified model can provide a powerful framework to link the local microscopic structure of active filaments to the large-scale collective phases that they form as a function of density.

**Figure 5:**
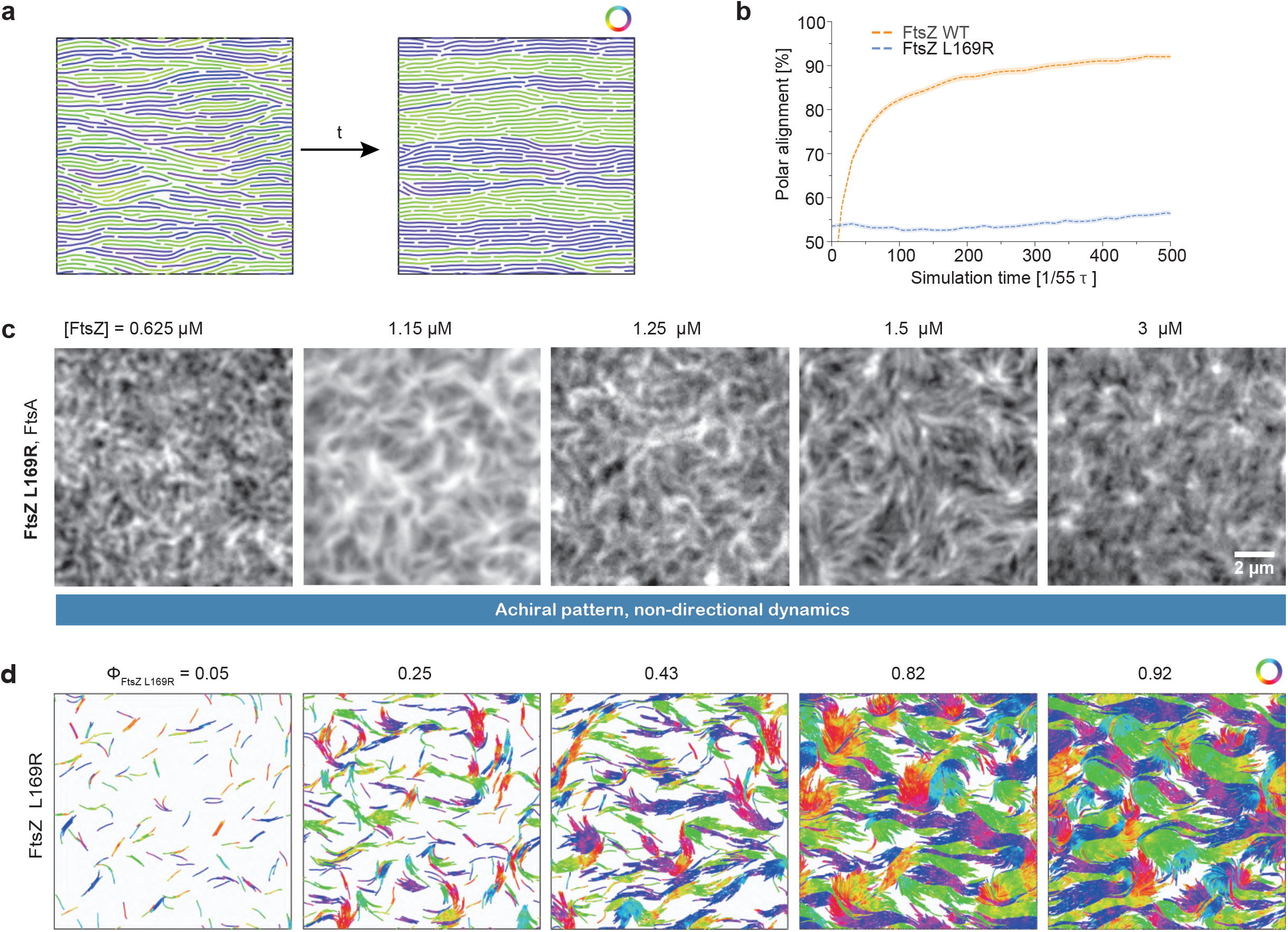
Microscopic sorting and large-scale organization of FtsZ L169R filaments. **a**, Toy simulations studying the polarity sorting kinetics of filaments with varying intrinsic curvature, length and bending rigidity - snapshot of the initial (left) and final configuration (right) of FtsZ L169R filaments in high density 6 = 0.88. Only bonds of the filaments (without the full diameter of beads) are presented for clarity. Filaments are color-coded according to the orientation of the bond vectors between beads. **b**, Fraction of parallel (polar) alignment of filaments as the function of simulation time. The FtsZ L169R was simulated with two times higher persistence and length than WT, no intrinsic curvature and 8x lower *Fp* (parameters *Pe =* 300, *ℱ =* 20, *kbend* = 107 *kBT/rad2, LfL169R* = 16*d*, /_0_ = 9, 6 = 0.22 - 0.88). While FtsZ WT prefers to align in a parallel orientation, FtsZ L169R stays aligned in nematic fashion until the end of simulations. **c**, Representative TIRF micrographs of Alexa488-FtsZ L169R at increasing FtsZ L169R and constant FtsA concentration. We observe very static thread-like self-organization of filaments with no concentration-dependent transitions. **d**, Snapshots of large-scale FtsZ L169R simulations with increasing density. The FtsZ L169R filaments were modeled with altered properties according to the HS-AFM analysis: filaments were 2x longer and more rigid, had no intrinsic curvature and 8x lower *Fp* resulting in 3x lower Peclet number and 2x lower flexure number than FtsZ WT. (parameters *Pe* = 300, *ℱ* = 20, *kbend* = 107 *kBT/rad2* /_0_ = 9, *LfL169R* = 16 *d)*. FtsZ L169R exhibits static thread-like pattern in all densities, as in the experiment. Filaments are color-coded according to the orientation of the bond vectors between beads.

## Discussion

In this study, we have identified density-driven transitions in the self-organization of FtsZ filaments between three phases: a low density disordered phase where individual filaments treadmill in a chiral manner with little interactions, an intermediate density phase where coherent chiral vortices form - as previously shown theoretically and experimentally^2,28,30^, and a high density phase where individual filaments straighten and where the system displays a uniform, nematic-like organization. Using a minimal model of active, chiral and flexible filaments we computationally analyzed how the physical properties of self-propelled filaments determine their large-scale self-organization. Although the presence of rings in intermediate densities can arise in the context of purely repulsive rigid filaments, a key finding from our computational work is that varying filament bending rigidity and attractive interactions changes qualitatively and quantitatively the phase diagram of possible morphologies. We find that the experimental system is consistent with intermediate values of both parameters, as filament flexibility in particular significantly modifies ring statistics and lifetimes. With semiflexible filaments, we observe indeed a competition between chiral shape, active treadmilling and inter-filament interactions, which at intermediate densities results in the co-existence and interconversion of chiral vortices and nematic-like traveling bands. This recapitulates well our experimental observations made on different spatial scales: while TIRF and STED microscopy allowed us to visualize the large-scale spatiotemporal dynamics of FtsZ filaments - and thus the transitions between different phases - HS-AFM revealed the behavior of individual filaments at different filament densities on a membrane surface. This data confirmed that the transition from chiral vortices to a nematic-like organization with increasing density goes along with straightening of the filaments, arguing for a relatively low FtsZ bending rigidity.

Furthermore, we demonstrated that a local perturbation, in this case a specific residue in the primary sequence of a protein, can dramatically change both individual filament properties and collective self-organization. Specifically, we found that the presence of an additional salt bridge between two monomers in filaments of FtsZ L169R increases their rigidity and lowers their intrinsic curvature. As a consequence, this mutant does not show a density dependent transition between different phases and instead always displays achiral, non-directional large-scale cytoskeletal networks. FtsZ L169R was originally described as bundling mutant that shows enhanced lateral interactions^38,39^. Our data suggests that enhanced lateral interactions are a consequence of mutant filaments being longer, more rigid and straighter and that the closer contact between them is only a secondary effect. The lack of a preference for a polar orientation of straight filaments are also in agreement with the observation of aberrant rings and spiral structures of FtsZ L169R found *in vivo*^38^. Although the intrinsic curvature of wildtype FtsZ filaments is much lower than that of MreB^43^ and FtsA filaments^44^, it could contribute to the correct alignment of the Z-ring perpendicular to the long cell axis in particular at early stages of cell division^45^.

Several experimental examples of chiral active matter have emerged in the past few years, across many different length scales. At the organismal scale, malaria parasites have been shown to have flexible rod-like shape and migrate actively in a chiral manner with different parasites having opposite chirality. This leads to sorting based on chirality, which is favored by mechanical flexibility^46^. Starfish embryos, although spherical in shapes, have also recently been shown to swim in a chiral manner, and collectively form crystal-like structures with odd elastic behavior^47^. At the cellular scale, chirality can bias the active nematic instabilities observed in confluent monolayers ^48^ and biofilms^49^. This has further been proposed to arise from cytoskeletal organization, due to the polar helicoidal structure of active filaments^50^, highlighting the need to a better understanding of the collective dynamics bridging different scales. In particular, it would be interesting in the future to investigate more complex models of filament treadmilling, for instance incorporating finite lifetimes or stress-dependent polymerization. Furthermore, our experimental and theoretical data suggests that weak and transient lateral interactions lacking any apparent biochemical specificity are sufficient for the alignment of treadmilling filaments on a membrane surface. Indeed, *in vivo* filaments in the Z-ring can move either in the same or opposite direction^51,52^ and recent experiments in Bacillus subtilis suggest that filament treadmilling facilitates their encounters promoting condensation of filaments into the Z-ring^23^. Relying on weak, non-specific interactions instead of specific residue contacts is therefore likely advantageous for the cell as it allows for the condensation of the Z-ring, while still permitting its dynamic reorganization. The relatively low bending rigidity that we observe for FtsZ could also be key to allow the Z-ring to adapt to the decreasing diameter of the constricting cell septum. Overall, our study highlights how minimal models of active matter based on symmetries can provide quantitative insights into fundamental biological functions.

## Supplementary Figure captions

**Supplementary Figure 1:**
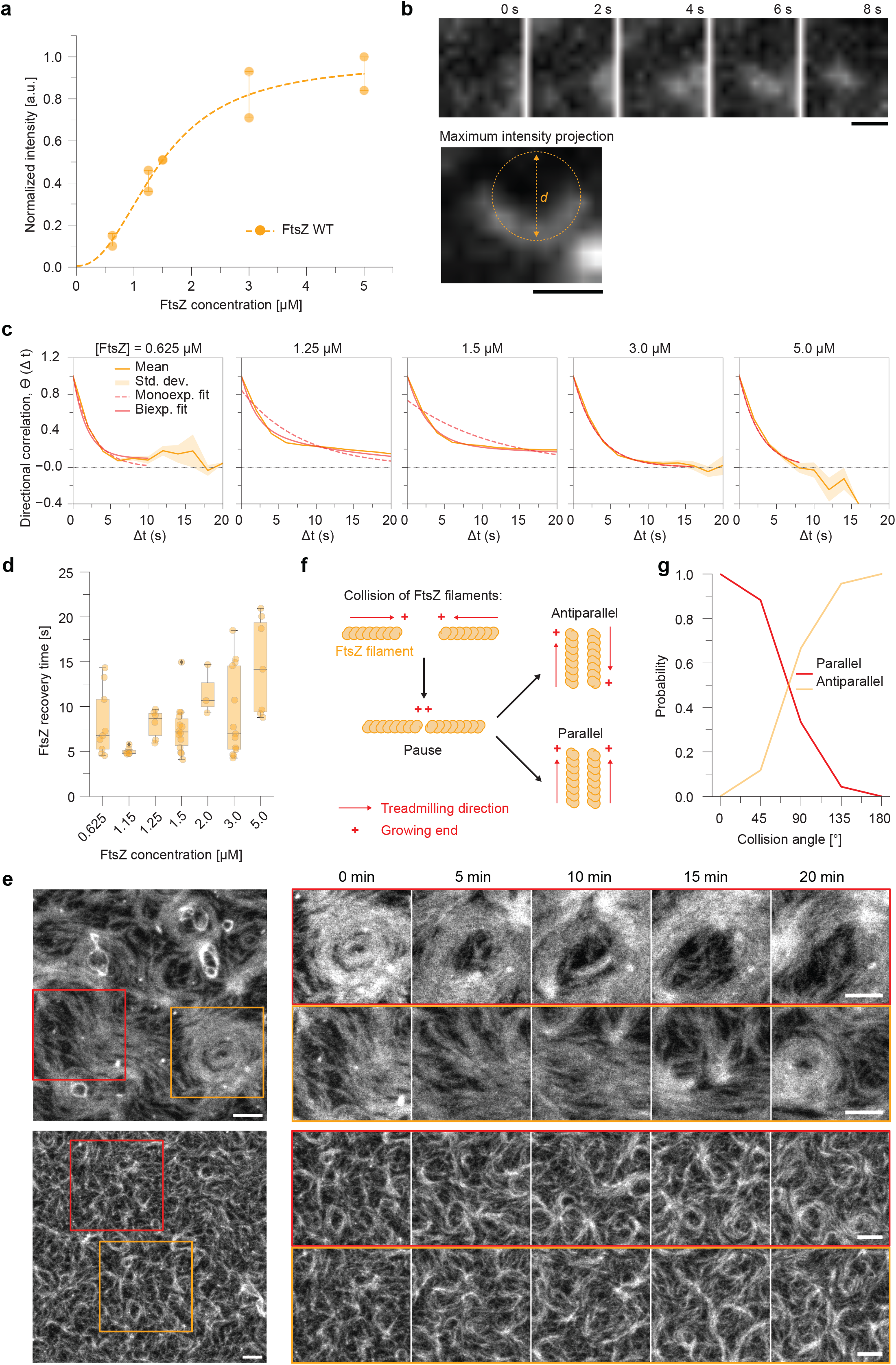
Characterization of FtsZ WT organization at increasing densities with TIRF and STED. **a**, Quantification of the intensity of Alexa488-FtsZ WT during a TIRF titration experiment. The density of FtsZ filaments on the supported membrane is saturated at ∼3µM. **b**, Snapshots of time lapse movie of a single FtsZ filament treadmilling on a membrane surface. The maximum intensity projection reveals the curved trajectory of the filament. **c**, Representative fits of mono-and bi-exponential functions to the directional autocorrelation of treadmilling trajectories. **d**, Quantification of the membrane residence time of FtsZ WT by FRAP experiments with increasing bulk concentrations. **e**, Representative micrographs of STED time-lapse experiments of Atto633-FtsZ WT. The insets show either rings which dynamically re-arrange into bundles (red) or bundles which re-arrange into rings (orange). The scale bars are 1µm. **f**, Model drawings showing the different types of collisions FtsZ filaments can undergo. In the aftermath of collision events FtsZ filaments align either parallel or antiparallel. **g**, Probability of parallel and antiparallel alignment as a function of impact angle.

**Supplementary Figure 2:**
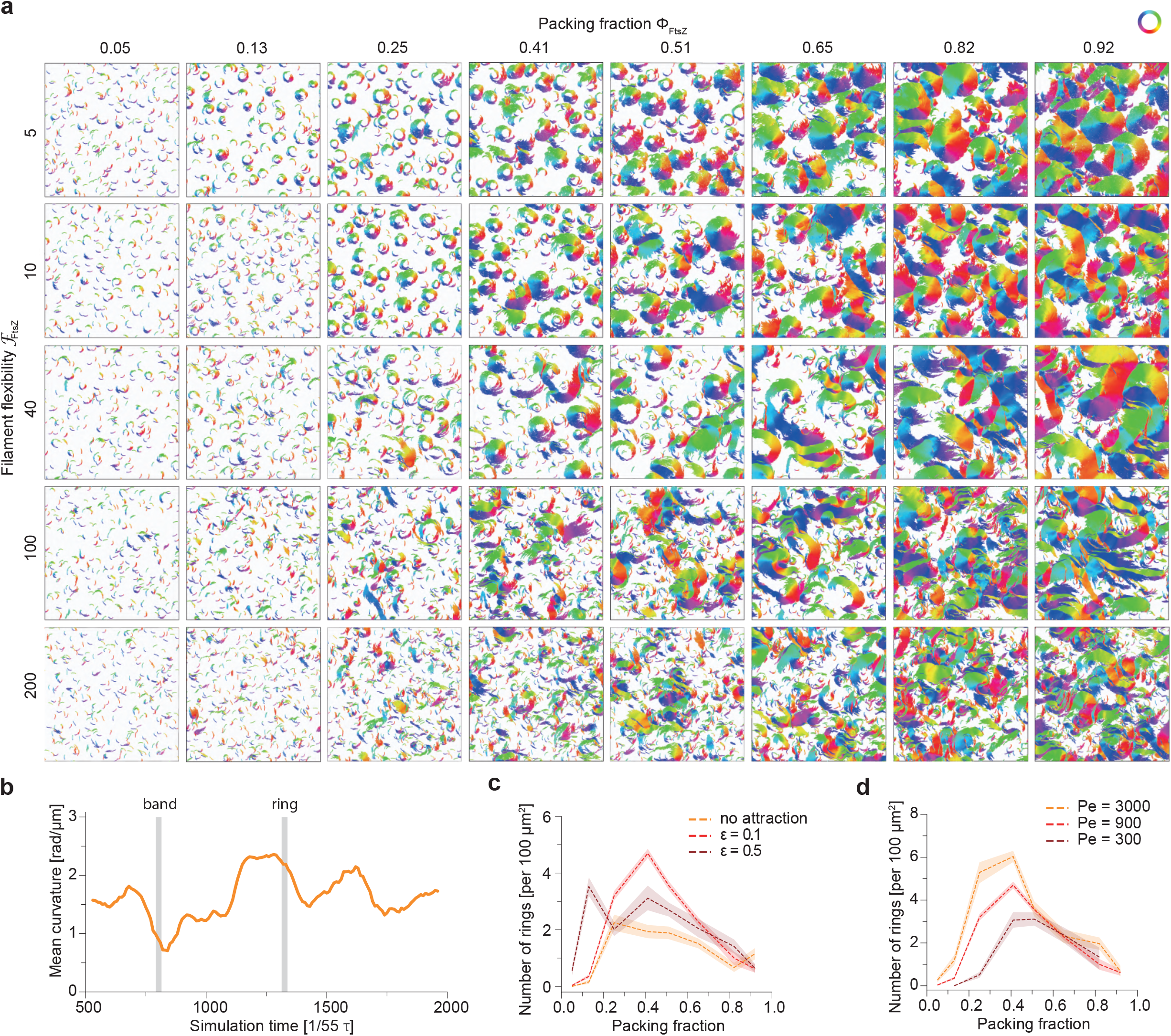
Numerical model of FtsZ WT - self-organization with varying rigidity, attraction and noise. **a**, Extended visual phase diagram of the large-scale patterns (*L* = 212 8) with varying filament flexibility (measured by flexure number *ℱ*, vertical axis) and density (horizontal axis). Filaments are colored according to the orientation of the bond vectors between beads. **b**, Evolution of average curvature of the system showing the temporal coexistence of chiral rings and polar bands in a small system size (*L* = 42 8) of intermediate density (*Φ* = 0.5). The vertical lines represent the timepoints of the snapshots in Figure 2c. **c**, Dependence of density of rings in the large-scale system (*L* = 212 8) on the packing fraction, for simulations with varying filament attraction. **d**, Dependence of density of rings in the large-scale system on the packing fraction, for simulations with varying Peclet number.

**Supplementary Figure 3:**
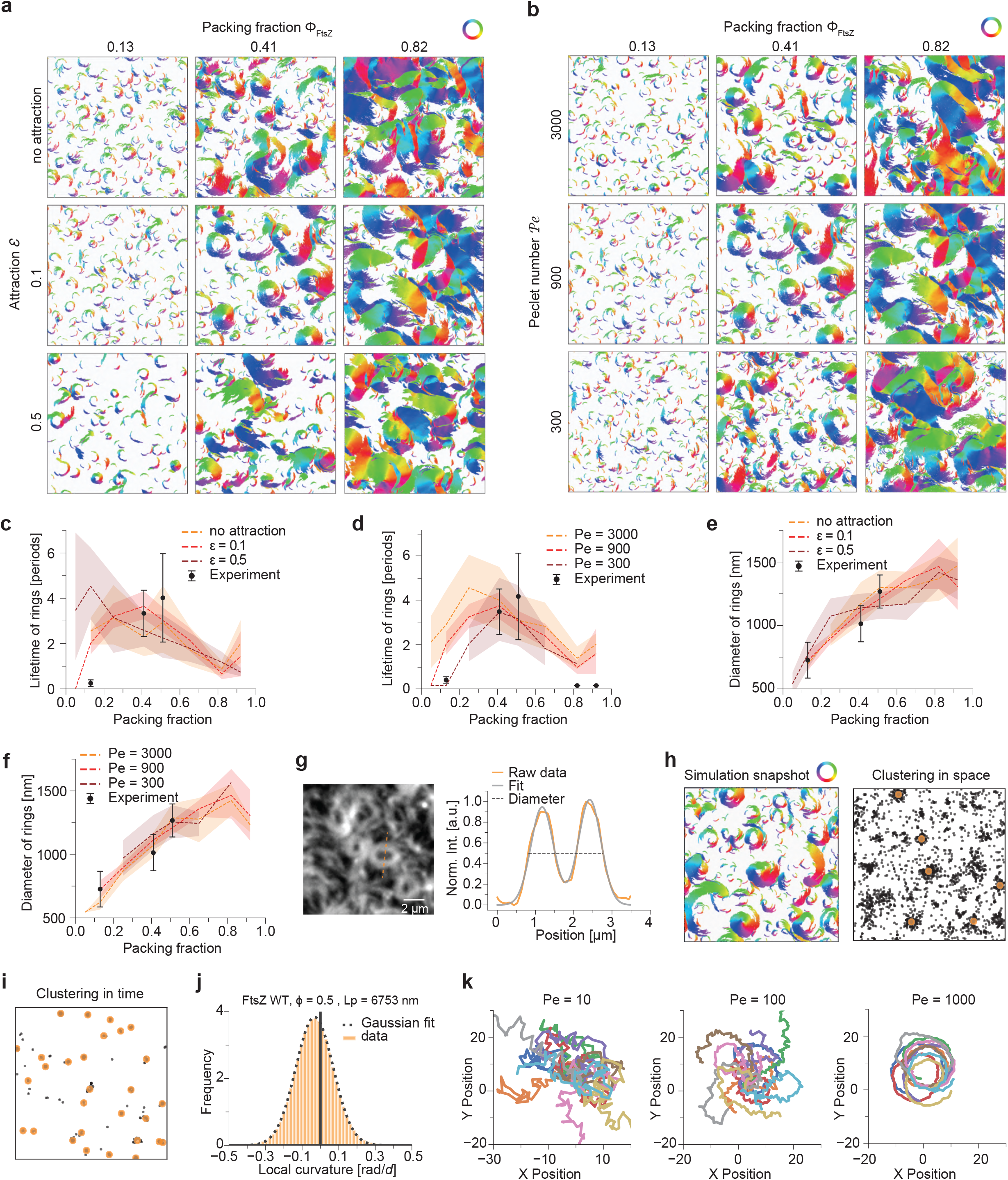
Numerical model of FtsZ WT - self-organization with varying attraction and noise, quantitative analysis. **a**, Visual phase diagram of the large-scale patterns (*L* = 212 *d*) with varying attraction (vertical axis) and density (horizontal axis). The rings are less stable without lateral attraction, while with the attraction being too strong, rings are favored in small density (*Φ* < 0.25), while in the intermediate density the cluster formation is very rapid resulting in lower occurrence of rings. Filaments are colored according to the orientation of the bond vectors between beads. **b**, Visual phase diagram of the large-scale patterns (*L* = 212 *d*) with varying noise (measured by Peclet number *Pe*, vertical axis) and density (horizontal axis). The rings are more abundant and stable with higher Peclet numbers with strongest differences appearing in lower density (0.05 < *Φ* < 0.25). Filaments are colored according to the orientation of the bond vectors between beads. **c**, Quantitative comparison of ring lifetime between large-scale simulations with varying filament attraction and experiments. The remaining simulation parameters were kept constant. **D**, Quantitative comparison of ring lifetime between large-scale simulations with varying Peclet number (noise) and experiments. The remaining simulation parameters were kept constant. **E**, Quantitative comparison of ring diameter between large-scale simulations with varying filament attraction and experiments. The remaining simulation parameters were kept constant. **F** Quantitative comparison of ring diameter between large-scale simulations with varying Peclet number (noise) and experiments. The remaining simulation parameters were kept constant. **H**, Example illustrating the automatic detection of rings. Left box depicts a simulation snapshot with filaments forming rings and bands (with filaments are colored according to the orientation of the bond vectors between beads), while the right box shows the result of clustering of filament rotation centers in space after applying all cutoffs (cluster size, polarity and effective radius). **i**, Result of the ring detection algorithm for the whole simulation after applying the time clustering algorithm to detect the rings in all calculated simulation snapshots. **j**, Analysis of filament persistence in the numerical simulations based on the distribution of local curvatures. Persistence length was extracted from variance of Gaussian function fitted to the data. **k**, Single filament trajectories (10 independent simulations) with varying Peclet number. The comparison to single-filament trajectories analyzed in **Fig. 1b** allowed us to set a lower bound to Peclet number in simulations *Pe* = 100 (see Methods), as below this value the intrinsic curvature of filament trajectories wouldn’t be detectable due to stochasticity of movement.

**Supplementary Figure 4:**
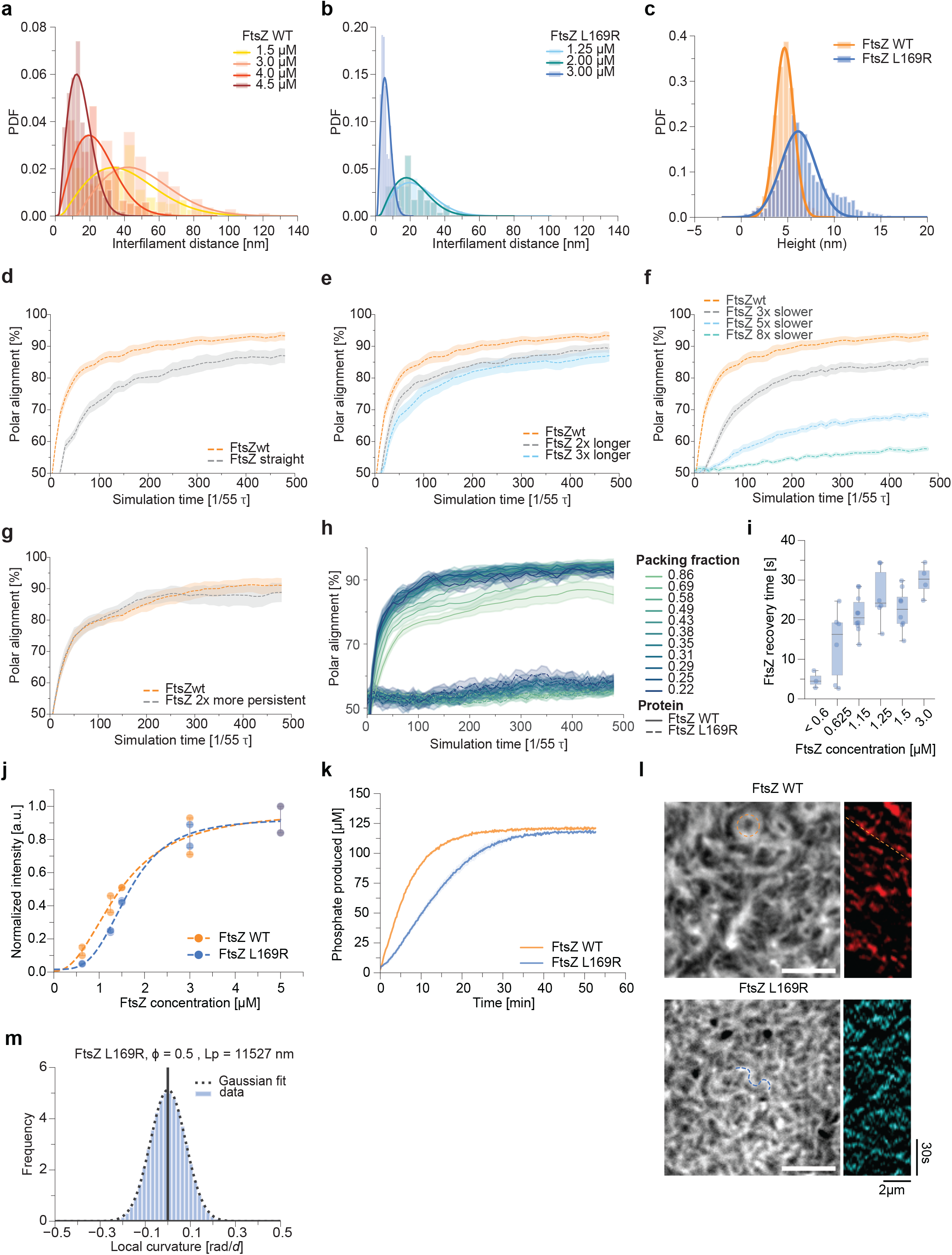
Characterization of FtsZ L169R and effect of filament properties on polarity sorting. **a**, The interfilament distance of FtsZ WT measured by HS-AFM. The distance decreases with increasing bulk concentrations and membrane densities of FtsZ WT. **b**, The interfilament distance of FtsZ L169R measured by HS-AFM. Filament distances are smaller than for the wildtype filaments even at lower FtsZ L169R concentrations. **c**, FtsZ WT (red) and L169R (blue) filaments shows no significant difference in their heights. **d**, Polarity alignment of simulations with FtsZ WT and FtsZ L169R in individual densities. The properties of mutant significantly slow down the polar alignment of filaments in all the studied packing fractions. The analyzed simulations correspond to the **Fig. 5b. e-h**, effect of individual filament properties on the polarity sorting. In each plot, simulations with FtsZ WT are presented to be compared with simulations of FtsZ WT with one changed parameter (intrinsic curvature, length, treadmilling speed, persistence length, respectively). All studied properties decrease the polar sorting with persistence length having the smallest effect. Each plot represents averaged results over four packing fractions in range to 6 = 0.29 to 6 = 0.88. **i**, Quantification of the membrane residence time of FtsZ L169R by FRAP experiments. Monomer turnover slows down with increasing FtsZ concentrations. **j**, Quantification of the TIRF intensities of Alexa488-FtsZ L169R (blue) at different protein concentrations. The intensity as well as the filament density saturates at ∼3µM, similar to FtsZ WT (red). **k**, Quantification of the GTPase hydrolysis rate of FtsZ WT (red) and FtsZ L169R (blue). The production of free phosphate is measured at a protein concentration of 5 µM. The rates are 2.2 ± 0.03 and 0.78 ± 0.03 GTP/FtsZ/min for WT and L169R respectively. **l**, TIRF micrographs and representative kymographs after differential imaging for FtsZ WT (left, red) and L169R (right, blue). The kymographs were obtained along the red or blue dashed lines. Diagonal lines in the kymograph of FtsZ WT correspond to directional polymerization dynamics of filament bundles, which are missing for L169R. Scale bars are 2 µm.

## Supplementary movies captions

**Supplementary movie 1: TIRF time-lapse movies of FtsZ WT**

TIRF time-lapse movies of Alexa488-FtsZ WT at different concentrations ([FtsA] = 0.2µM). With increasing FtsZ concentration, the pattern changes from rotating rings and directional moving filament bundles to a more nematic pattern. Movies were acquired at 0.5 frame per second and correspond to **Fig. 1a**.

**Supplementary movie 2: STED time-lapse experiments of FtsZ WT**

STED time-lapse movies of Atto633-FtsZ WT. The FtsZ concentration was 1.25µM and FtsA 0.2µM. The experiments were recorded at 1 frame every 4-6s, where the acquisition rate depends on the field of view. The movies display the co-existence of rings and bundles of FtsZ filaments. The movies correspond to **Supplementary Fig. 1c**.

**Supplementary movie 3: FtsZ phase diagram**

Large scale FtsZ patterns (L = 212 *d*) with varying filament flexibility (measured by flexure number *ℱ*, vertical axis) and packing fractions (horizontal axis). Filaments are colored according to the orientation of the bond vectors between beads. We observe ring-like self-organization of rigid filaments (*ℱ* = 5), spatial coexistence of chiral rings and polar bands in regime of semiflexible filaments (*ℱ* = 40) and disordered patterns with flexible filaments (*ℱ* = 200). Movies correspond to **Fig. 2b**.

**Supplementary movie 4: Temporal coexistence of rings and bands**

Temporal coexistence of chiral rings and polar bands in a small system (L = 42 *d*) of intermediate density (6 = 0.5) and filament flexibility (*ℱ* = 40). With increasing density, the ring state becomes unstable and filaments organize only in bands. Filaments are colored according to the orientation of the bond vectors between beads. Movie corresponds to **Fig. 2c** and **Supplementary Fig. 2b**.

**Supplementary movie 5: FtsZ topological defects**

Topological defects in high filament density (6 =0.9) and small system size (L = 42 *d*). The rigid filaments (*ℱ* = 5) form spiral (+1) topological defects, whereas semiflexible filaments (*ℱ* = 40) form only nematic defects (+1/2 and -1/2) due to the filament straightening. Only bonds of the filaments (without the full diameter of beads) are presented for clarity. Filaments are colored according to the orientation of the bond vectors between beads. Movies correspond to **Fig. 2e,f**.

**Supplementary movie 6: HS-AFM time-lapse movies of FtsZ WT**

HS-AFM movies of FtsZ WT filaments. As the density of FtsZ WT filaments on the supported lipid bilayer increases, they become less dynamic and straighter. At high densities, the filaments show a nematic order with topological defects. Time-lapse movies were acquired with 3 and 2 frames per second. The movies correspond to **Fig. 3a, 3d**.

**Supplementary movie 7: HS-AFM time-lapse experiments of FtsZ L169R**

HS-AFM videos of FtsZ L169R filaments. As the density of FtsZ L169R increases, but the filaments remain straight and static. At high densities, the filaments pack extremely tight together. Images were acquired with 3 and 2 frames per second. The movies correspond to **Fig. 4b**.

**Supplementary movie 8: Polar sorting of filaments with different properties**

Polar sorting of filaments with properties of FtsZ WT and FtsZ L169R in two different densities. FtsZ L169R is 2x longer and more rigid, non-chiral and has lower self-propulsion than FtsZ WT, resulting in 3x lower Peclet number and 2x lower flexure number. Only bonds of the filaments (without the full diameter of beads) are presented for clarity. Filaments are colored according to the orientation of the bond vectors between beads.

**Supplementary movie 9: TIRF time-lapse movies of FtsZ L169R**

TIRF time-lapse movies of Alexa488-FtsZ L169R at increasing concentrations ([FtsA] = 0.2µM). FtsZ L169R does not form rings as seen for FtsZ WT and the pattern appears less dynamic. Movies were acquired at 0.5 frame per second and correspond to **Fig. 5c**.

**Supplementary movie 10: Large-scale simulations of FtsZ L169R**

Large-scale simulations with increasing concentration of FtsZ L169R. The FtsZ L169R filaments are non-chiral, 2x longer and more rigid than FtsZ WT and are self-propelled with 8x lower speed, resulting in *Pe* = 200 and *ℱ* = 20 and filament persistence length 2x longer than FtsZ WT. FtsZ L169R does not form rings and self-organizes into less dynamic pattern. Filaments are colored according to the orientation of the bond vectors between beads. Movies correspond to **Fig. 5d**.

## Material & Methods

### Experiments

#### Protein biochemistry

Proteins used in this study, wildtype FtsZ, FtsZ L169R and FtsA, were purified as previously described^53^. FtsZ, L169R was obtained by site-directed mutagenesis (SDM). Leucine 169 was replaced with Arginine, by exchanging two nucleotides (CTG → CGC). FtsZ L169R was purified as the wild-type protein as described before for the wild-type protein.

#### Preparation of coverslips

We used piranha solution (30 % H2O2 mixed with concentrated H2SO4 at a 1:3 ratio to clean the glass coverslips for 60 min. This was followed by extensive washes with double-distilled H2O, 10 min sonication in ddH2O and again washing in ddH2O. The coverslips were used within one week and were stored in ddH2O water. Furthermore, before coverslips were used to form supported lipid bilayers, they were dried with compressed air and treated for 10 min with a Zepto plasma cleaner (Diener electronics) at maximum power. As reaction chambers we used 0.5 ml Eppendorf tubes missing the conical end, which were glued on the coverslips with UV glue (Norland Optical Adhesive 63) and exposed to ultraviolet light for 10 min.

#### Preparation of small unilamellar vesicles (SUVs)

DOPC (1,2-dioleoyl-sn-glycero-3-phosphocholine) and DOPG (1,2-dioleoyl-sn-glycero-3-phospho-(1’-rac-glycerol)), which were purchased from Avanti Polar Lipids, at a ratio of 67:33 mol% were used. The lipids in chloroform solution were mixed inside a glass vial in the appropriate volumes and dried with filtered N2 for a thin lipid film. Remaining solvent was removed by putting the lipids in a vacuum desiccator for 2 h. Afterwards swelling buffer (50 mM Tris-HCl [pH 7.4] and 300 mM KCl) was added to the lipid film to obtain a lipid concentration of 5 mM. After incubating the suspension for 30 min at room temperature, the multilamellar vesicles were vortexed rigorously and freeze–thawed (8x) in dry ice or liquid N2. The liposomes were tip-sonicated using a Q700 Sonicator equipped with a ½ mm tip (amplitude = 1, 1 second on, 4 seconds off) for 25 min on ice to obtain SUVs. Finally, the vesicles were centrifuged for 5 min at 10,000 *g* and the supernatant was stored at 4 °C in an Argon atmosphere and used within one week.

#### Preparation of supported lipid bilayers (SLB) for TIRF

SLBs were prepared by diluting the SUV suspension to a concentration of 0.5 mM with reaction buffer supplemented with 5 mM CaCl2. SLBs were incubated for 30 min at 37 °C and non-fused vesicles were washed away by 8×200 µL washes with reaction buffer (50 mM Tris-HCl [pH 7.4], 150 mM KCl and 5 mM MgCl2). The membranes were used within 4 hours.

#### Total internal reflection fluorescence (TIRF) microscopy

Experiments were performed using a Visitron iLAS2 TIRF microscope, equipped with a 100xOlympus TIRF NA 1.46 oil objective. The fluorophore Alexa488 was excited with a laser line at 488nm. The emitted fluorescence from the sample was filtered using a Laser Quad Band Filter (405/488/561/640 nm). A Cairn TwinCam camera splitter equipped with a spectral long pass of 565 nm and a band pass filter of 525/50 nm was used. Time series were recorded using Photometrics Evolve 512 EMCCD (512 × 512 pixels, 16 × 16 μm^2^) operating at a frequency of 5 Hz.

#### Stimulated emission depletion (STED) microscopy

STED microscopy was performed at room temperature on an inverted Expert Line STED microscope (Abberior Instruments) with pulsed excitation and STED lasers. A 640 nm laser was used for excitation and a 775 nm laser for stimulated emission. A oil immersion objective with 1.4 NA (Olympus, UPLSAPO 100XO) was used for image acquisition. The fluorescence signal was collected in a confocal arrangement with a pinhole size of 0.8 airy units. For detection a 685/70 nm bandpass filter (Chroma, #F49-686) and a photon counting avalanche photodiode (Laser Components, Count-T100) were used. The pulse repetition rate was 40 MHz and fluorescence detection was time-gated. Data were acquired with 10 μs pixel dwell time and 30 nm pixel size for time lapse imaging and 20 µs with 20 nm pixel size for overview images, 5 – 6.5 µW excitation laser power and 30 – 40 mW STED laser power. The power values refer to the power at the sample, measured with a slide powermeter head (Thorlabs, S170C). A spatial light modulator (SLM) imprinted the STED phase pattern for lateral resolution increase. Image acquisition and microscope control were performed with Imspector software version 14.0.3052.

#### High speed atomic force microscopy (HS-AFM)

A laboratory-built tapping mode (2 nm free amplitude, ∼2.2 MHz) high-speed atomic force microscope (HS-AFM) equipped with a wide-range scanner (6µm x 6µm) was used to visualize the dynamics of the system. BL-AC10DS-A2 (Olympus) cantilevers were used as HS-AFM scanning probes. The cantilever has a spring constant (k) of 0.1N/m and a resonance frequency (f) of 0.6MHz in water or 1.5MHz in air. The dimensions of the cantilever are: 9 µm (length), 2 µm (width), and 0.13 µm (thickness). To achieve high imaging resolution, a sharpened and long carbon tip with low apical radius was made on the existing tip of the cantilever using electron-beam deposition (EBD) as described previously^54–56^. Scanning speed varied from 0.2s to 5s per frame. The number of pixels acquired were adjusted for every measurement depending on the scan size (min: 2nm, max: about 50nm). The in-house designed program “Kodec” was used to read the data generated by HS-AFM. The ware stores all parameters, calibration and description given during the measurement and allows to load a whole folder or several movies.

#### FtsZ TIRF and STED experiments on SLBs

To study the organization of increasing concentrations of treadmilling FtsZ filaments on supported lipid bilayers, we used 0.2 μM FtsA and increasing concentrations of Alexa488-FtsZ wt or L169R (0.625/1.25/1.5/3/5 μM) in 100 µl of reaction buffer. Additionally, the reaction chamber contained 4 mM ATP/GTP and a scavenging system to minimize photobleaching effects: 30 mM d-glucose, 0.050 mg ml^−1^ Glucose Oxidase, 0.016 mg ml^−1^ Catalase, 1 mM DTT and 1 mM Trolox. Prior addition of all components a corresponding buffer volume was removed from the chamber to obtain a total reaction volume of 100 µl. The FtsZ filaments was imaged by TIRF at one frame per two seconds and 50 ms exposure time.

STED microscopy exposes fluorescent labelled proteins to a much higher laser intensity compared to TIRF and imposes bleaching and photo-toxic effects on the FtsZ filaments. We adjusted our imaging setup accordingly to avoid bleaching and photo-induced bundling of FtsZ filaments. First we replaced Alexa488 with Atto633, a widely used STED dye with higher photon yield and improved resistance to bleaching. Furthermore, we optimized the scavenging solution used for imaging: 60 mM d-glucose, 0.10 mg ml^−1^ Glucose Oxidase, 0.032 mg ml^−1^ Catalase, 20 mM DTT and 2 mM Trolox. Finally, we changed the acquisition protocol to pixel-step based excitation/STED cycles. Here, we introduce short breaks in between the excitation cycles as follows: 5µs excitation/STED - 10µs break - 5µs excitation/STED - 10µs break^57^. Together, these changes allowed us to observe the same behavior as in TIRF microscopy experiments, but with increased resolution for longer than 10 minutes. For the experiments shown, we used 0.2 µM FtsA and 1.5µM Atto633-FtsZ. In the time-lapse montages shown in S1 the FtsZ pattern was imaged at one frame per five seconds.

#### Preparation of SLBs for HS-AFM

SUVs were prepared as described above. An ultra-flat muscovite mica layers (1.5 mm Ø) substrate was mounted on a glass stage using a standard 2-component glue. The glass stage was then attached to the scanner with a thin film of nail polish. A drop of acetone was deposited on the stage/scanner interface to ensure a flat nail polish layer. The mounted stage was dried at RT for about 30 min. A fresh cleaved mica layer was used as substrate to form a supported lipid bilayer (SLB) by depositing ∼4 µL of a mix of 1 mM SUVs suspension in reaction buffer with additional 5mM CaCl2. To avoid drop breakage, the scanner was flipped upside down and inserted in a custom-made mini-chamber with a thin water film at the bottom (a 500 µL tube cut on the bottom and glued to a petri dish). The drop was incubated on the stage for at least 30 min. After, the drop was exchanged 5-10 times with 5µL of fresh reaction buffer. The stage was immediately inserted in the HS-AFM chamber containing about 80µL of the same reaction buffer. Prior to the addition of the proteins, HS-AFM imaging and indentation were performed to assess the quality of the SLB. When the force-distance curve showed the typical lipid bilayer indentation profile (∼2-4 nm) the SLB was used in the next steps.

#### FtsZ HS-AFM experiments on SLBs

The selected proteins (FtsZ WT or L169R, FtsA) were added to the chamber with ATP/GTP (4mM each) and DTT (1mM) (final concentrations). Optimal protein concentration and ratios were tested as well for a range of ∼0.3-4.5 µM for FtsZ and 0.05-1.2 µM FtsA.

### Image processing and analysis

For data analysis of TIRF and STED experiments, the movies were imported to the ImageJ Version 2.9.0/1.53t ^58^ software and raw, unprocessed time lapse videos were used. The contrast of micrographs in the manuscript was adjusted to improve visibility of filaments. HS-AFM data was exported to ImageJ, where the post-processing, such as noise reduction and smoothing, was carried out. Noise reduction and smoothing were performed using a bandpass filter of different size depending on the image features. Analysis of HS-AFM movies was performed in MatLab, using the AFM analysis package FiberApp^59^ (Version 2017b of MatLab). HS-AFM micrographs in the manuscript are raw data, where the contrast was optimized for best quality.

#### FtsZ intensity analysis

To estimate the saturating coverage of FtsZ on SLBs we titrated bulk FtsZ concentrations from 0.625 – 5 µM in two independent experiments on the same day, to avoid any changes in the power of the microscope. After reaching equilibrium (20 minutes), we measured the intensity at three different field of views at each concentration. Finally, we normalized these intensity values by a min-max normalization and fitted a Hill 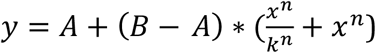, where A is the starting point, B the ending point and n the Hill coefficient (**Supplementary Fig. 1a** and **Supplementary Fig. 3j)**.

#### FtsZ trajectory analysis

To estimate the diameter of trajectories of FtsZ single filaments, we performed a maximum intensity projection of selected regions of interest (ROI) showing isolated single filaments. Next, a circle was manually drawn on top of the filament trajectory and its diameter measured (**Fig. 1b)**.

#### Differential imaging and directional autocorrelation

To quantify at the directional flows of the FtsZ filament pattern, we used a previously developed automated image analysis protocol^35^. In this routine, we first track the growing ends of FtsZ filament bundles and then compute the directional autocorrelation from these trajectories. The corresponding correlation coefficient is obtained by computing the correlation of the angle between two consecutive displacements as a function of an increasing time interval (*∆t*) and is a measure for the local directional persistence of treadmilling trajectory *i*. The autocorrelation curves were best fitted assuming a fast and slow decay, whose rates were extracted by fitting a two-phase exponential decay y = *a*1 ∗ *e*^(−*b*1∗*t*)^ + *a*2 ∗ *e*^(−*b*2∗*t*)^, where a1, a2 are the starting points and b1, b2 are the fast and slow decay rates. The two half times of the two-phase exponential decay were calculated from the respective decay rates (**Fig. 1c** and **Supplementary Fig. 1c**).

#### Fluorescence recovery after photo-bleaching (FRAP) analysis

For FRAP experiments, a small area of the membrane was bleached with a high 488nm laser intensity after the pattern has reached equilibrium. To obtain the recovery half-time we used a Jython macro script for ImageJ (Image Processing School 8 Pilsen 2009) to fit the fluorescence recovery with *I*(*t*) = *a*(1 − *e*^*−bt*^), where I(t) is the intensity value corrected for photobleached effects. FRAP experiments were acquired with an exposure time of 50msec and an acquisition time of one frame every 1 second (**Supplementary Fig. 1d** and **Supplementary Fig. 3i**).

#### Collision angle analysis from STED movies

The angle of FtsZ filaments collision was measured using the Angle tool of ImageJ. Three points, corresponding to the incoming FtsZ and the center point of the collision, respectively, were manually set. The subsequent alignment (parallel or antiparallel) was evaluated manually (**Supplementary Fig. 1g**).

#### Quantification of GTP hydrolysis rate of FtsZ

To measure the GTPase rate of FtsZ WT and L169R we used the commercially available EnzCheck kit (ThermoFisher, E6646). The protein was buffer exchanged into phosphate free SLB reaction buffer (150mM KCl, 50mM Tris, 5mM MgCl2, pH 7.4). The proteins were then diluted to 5µM together with the 20x reaction buffer of the kit, the SLB reaction buffer, MSG and PNP. The reaction was incubated for 15 minutes to remove any traces of free phosphate with the PNP. Subsequently 200µM GTP were added and the production of free phosphate by FtsZ was measured. The appropriate controls (Buffer + GTP, Buffer + FtsZ WT/L169R) were measured simultaneously and subtracted from the Buffer + FtsZ WT/L169R + GTP curves. The rates are extracted from the slope of the initial linear increase from 0-5 minutes (**Supplementary Fig. 4k**).

### Ring analysis

#### Ring density

the number of rings in a given field of view were manually counted after averaging every 40 seconds of a time lapse movie **(Fig. 2g)**.

#### Ring diameter and width

First, to remove imaging noise fast intensity fluctuations, we first calculated a walking average over five frames of a time-lapse movie. Next, we obtained the fluorescence intensity profile across the ring diameter using and further analyzed this profile by background intensity subtraction and fitting a double Gaussian function with the Scipy module [2]. From this fit, the ring width can be obtained as the full width at half maximum (FWHM) averaged over the two peaks. The outer and inner diameters were computed as distances between the corresponding FWHM values. The outer diameter of rings was reported and compared to simulations **(Fig. 2i, Supplementary Fig. 3g)**.

#### Ring lifetimes

were obtained by first preparing a kymograph along the diameter of a ring and then measuring the length of two parallel vertical lines corresponding to the lifetime or the ring **(Fig. 2h)**.

### Single filament analysis

HS-AFM movies were imported to FiberApp^59^ and FtsZ filaments were traced. To simplify the tracking of FtsZ filaments, we used the A* pathfinding algorithm with 100 iterations. Furthermore, we used the open contour type, and the parameters were: Alpha = 10, Beta = 10, Gamma = 20, Kappa1 = 20, Kappa2 = 10. The fiber intensity was measured directly on the analyzed frame. After fitting all filaments within one frame, we extracted the persistence length (Lp) from calculating the mean-squared end-to-end distance :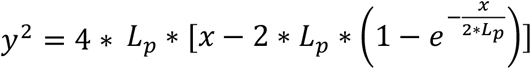. The contour length was extracted by measuring the average length distribution of all frames within a specific FtsZ density. The local curvature was extracted directly from the xy-coordinates of the filaments, obtained from Fiber app. Subsequently, we averaged the local curvatures of each filament to obtain one curvature value per filament at different densities (**Fig. 3c, 4c-4e**).

### Numerical simulations

The simulations of FtsZ filaments are based on the self-propelled worm-like chain model^31^, extended for polymer chirality (**Fig. 2a**). A filament is represented by *N+1* beads with radius *a0* connected by *N* stiff bonds and chiral bending potentials. To minimize friction between the polymers the bond length is chosen to be equal to *a0*, leading to overlaps of neighboring beads and filament length *Lf = N a0*. The overdamped equation of motion is therefore given by:

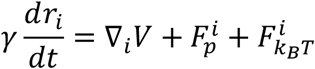

where *ri* are the coordinates of the beads, γ is the friction coefficient, 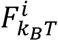 is the thermal noise force (modeled as white noise with zero mean and variance *4kBTγ/δt*, where *δt* is the simulation timestep and *kBT* is the effective temperature), and *V* is the potential energy, which comprises of both intrafilament and inter-filament interactions:

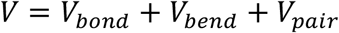

where the first term is a harmonic bond potential penalizing filament stretching

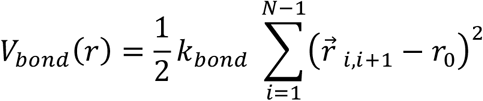

while the second term is a harmonic bending potential penalizing filament bending

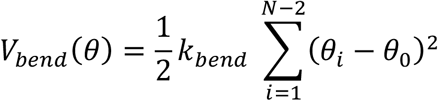

where 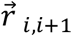 is the bond vector between the neighboring beads, *θ*_*i*_ is the angle between the neighboring bonds, *r0* and *θ*_0_ represent the equilibrium bond length and rest angle (with *kbond* and *kbend* the corresponding spring constants). The existence of a non-zero rest angle implies an intrinsic and chiral curvature for individual filaments, while the *kbond* parameter is used large enough to keep the bond length in the polymer constant. Finally, filaments interact via soft Lennard-Jones potential, which accounts for excluded volume interactions and potentially a middle range attraction

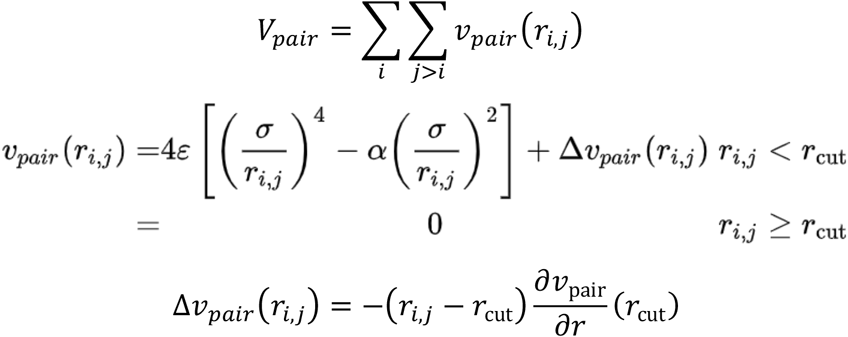

where *ri,j* is the vector between positions of beads *i and j* (which can belong to any filament), *ε* is the depth of the potential well, *σ* is distance where *Vpair* is zero and *rcut* is the cutoff distance of the potential. The potential is shifted by the subtraction of the value of the force at *rcut*, such that the force smoothly goes to zero at the cut-off. When accounting only for repulsive interactions, we use *rcut* consistent with the interaction minimum 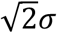. 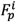 is the active self-propulsion force that mimics the filament treadmilling (acting tangentially along the bonds of the polymer):

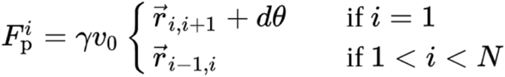

where *dθ* = *π* − *θ*_0_ ensures the chiral self-propulsion of the first bead. The simulations are performed in 2 dimensions and employ periodic boundary conditions.

#### Details of the simulation setup & parameter exploration

HOOMD-blue v2.9^60^ was used to run the simulations, with in-house modifications of the chiral worm-like chain model. Specifically, to ensure the chirality of the polymers, an asymmetric bending potential was used, having a signed curvature (always calculated in the direction from head to tail of the filament).

Simulation parameters and results are reported in dimensionless form, where length is measured in units of the effective bead diameter *d* (defined as the interaction minimum 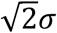), energies in units of the thermal energy *kBT*, and time in units of the filament rotation periods v. The equations of motion were numerically integrated using the Euler scheme with timestep *δt* = 1.8 • 10^−5^*τ*. The dynamics of filaments was governed mainly by two dimensionless numbers, namely the Peclet number

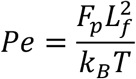

and flexure number

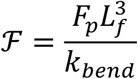

Unless stated otherwise, the simulation parameters to model FtsZ WT were: *Pe* = 900, *ℱ* = 40, *kbond* = 1000 *kBT/d*2, *γ = 1, kbend* = 53.5 *kBT/rad2*, bending angle *θ*_0_ = 3.08 rad, effective temperature for the thermal noise force 0.215 *kBT* and *rcut* = 1.7*d*. Since the directional motion of filament bundles are consistent with microscopic polar ordering of filaments (**Fig. 1d**), we modelled filaments with length *Lf* = 8 *d*, with aspect ratio corresponding to bundles with thickness of 5 FtsZ filaments^31^ To compare with experiment the reduced simulation units were recalculated with constants 1*d* = 50nm (corresponding to filament length FtsZ WT *Lf* = 400 *µm* and filament curvature 3.48 *rad/µm) and τ* = 78.5 *s* (considering a ring with diameter 1 5m and treadmilling speed 0.04 *µm/s*). We define packing fraction as *ϕ = NNfdr0/L2*, where *Nf* refers to the number of filaments and *L* to the box size. The simulated filament densities ranged from *ϕ* = 0.05 to *ϕ* = 0.9. The parameter space was explored by varying *Nf, kbend*, s*ε* and effective temperature *kBT* to alter filament density, flexure number, attraction and Peclet number, respectively.

To check if our explored parameter space is reasonable, we performed a few sanity checks. We computed the persistence length of filaments in our simulations from the distribution of local filament curvatures in intermediate density57. *Lp* was obtained by fitting a Gaussian function to local curvature data and extracting its variance: 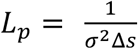, where *σ* is the standard deviation of Gaussian function and *Δ*Ñ is the filament contour spacing61. The resulting values *Lp* = 6753 nm for FtsZ WT (**Supplementary Fig. 3j**) and *Lp* = 11527 nm for FtsZ L169R (**Supplementary Fig. 4m**) agrees well with our HS-AFM analysis **(Fig. 4d)**. We also checked the order of magnitude of Peclet numbers of FtsZ WT in our simulations (**Supplementary Fig. 3k**) by comparing the single-filament trajectories to experimental trajectories from TIRF data (**Fig. 1b**).

The simulations of active filaments were initialized in a nematic configuration (each filament was placed on the lattice in a straight configuration randomly oriented either left or right). The initial time needed for the equilibration of the system (4v) is discarded in the analysis. Parameter screening and subsequent simulations, which were focused on the molecular detail (temporal interconversion of rings and bands in **Fig. 2c**, topological defects in **Fig. 2e,f**, polarity sorting in **Fig. 4a,b**) were performed with a box size *L* = 42 *d*, whereas all the reported large-scale patterns were simulated in the box of size *L* = 212 *d*.

To get a well-mixed state of non-chiral filaments with lower *Fp* (**Fig. 4a,d**), the system was initiated in low density (*ϕ* = 0.25 and lower) on a lattice in a nematic configuration with high thermal noise force (*Pe* = 100). The denser systems were initialized with enlarged boxes of density *ϕ* = 0.25, and after mixing of the filaments, these systems were down-scaled and equilibrated in a stepwise manner, to reach a high-density system with effectively random initil conditions.

The large-scale simulations (**Fig. 2b, 4d**) were run for 20 *τ*, whereas the small systems, where temporal interconversion of individual rings and bands was quantified (**Fig. 2c**), were run for 40 *τ*. Finally, polarity sorting simulations (**Fig. 4a,b**) were run for 10 *τ* and analyzed from the initial timestep. Each combination of parameters was run in at least 10 repeats for small system sizes and 5 repeats for larger systems.

#### Simulation analysis

Unless stated otherwise, simulation frames were analyzed with frequency of 0.2 *τ* after the equilibration. The average filament curvature (**Fig. 2d, 3b**) was computed by first calculating the mean curvature of each filament and then averaging it over a single snapshot of the system. The resulting distribution of the average curvatures consists of pooled data over both multiple simulation time points and simulation repeats.

To automatically detect rings in the system the instantaneous centers of rotation were computed for each filament in each analyzed snapshot. These points were subsequently clustered using Freud library^39,58^ with distance threshold related to the density of the system (cutoff defined as half of average distance between filaments: 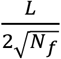. Clusters with more than 10 members and normalized radius (given by radius of gyration / cluster size) lower than 0.1 *d* were considered to be ring centers. To filter out unclosed rings, the normalized polarity (given by norm of the vector summing the filament orientations / cluster size) of filaments belonging to a ring was computed and had to be lower than 0.25 *d* (**Supplementary Fig. 3h**). Since stable rings didn’t show significant overall translation, we performed additional clustering of ring centers in time with distance cutoff 3*d* to track rings in time and get a lifetime of the rings (estimated by subtraction of last and first snapshots of each observed ring, **Fig. 2h, Supplementary Fig. 3c,d**). This clustering was also used to remove falsely detected “rings” by removing all clusters of size lower than 3 members (**Supplementary Fig. 3i**). All the cutoffs of this analysis were chosen and validated by visual comparison to simulation movies in all studied densities.

To calculate the average density of rings in simulations the number of rings in a snapshot was divided by the total area (**Fig. 2g, Supplementary Fig, 2c,d**). The diameter of rings was estimated by calculating the average distance of filaments from the center of the ring (**Fig, 2i, Supplementary Fig, 3e,f**).

The polarity alignment (**Fig. 5b, Supplementary Fig. 4d-h**) was analyzed by computing all the neighboring beads (not belonging to the same filament, with the same distance threshold *rcut* used in the pair potential) in the system using Freud library and classifying the neighbor relative orientation as either parallel or antiparallel based on their angle (< 90 degrees = parallel, > 90 degrees = anti-parallel).

#### Protein structure prediction

Protein modeling was performed with AlphaFold-Multimer v2^41,63^ implemented in Google Colab using the filament structure of Staphylococcus aureus FtsZ PDB 3VOB as template.

## Acknowledgements

This work was supported by the European Research Council through grant ERC 2015-StG-679239 by the Austrian Science Fund (FWF) StandAlone P34607 to M.L., and by the Kanazawa University WPI-NanoLSI Bio-SPM collaborative research program. Z.D. has received funding from Doctoral Programme of the Austrian Academy of Sciences (OeAW): [Grant agreement 26360]. We thank Jan Brugues (MPI CBG, Dresden, Germany), Andela Saric (ISTA, Klosterneuburg, Austria), Daniel Pearce (Uni Geneva, Switzerland) for valuable scientific input and comments on the manuscript. We are also thankful for the support by the Scientific Service Units (SSU) of IST Austria through resources provided by the Imaging and Optics Facility (IOF) and the Lab Support Facility (LSF).

